# Pathogenic tau accelerates aging-associated activation of transposable elements in the mouse central nervous system

**DOI:** 10.1101/2021.02.25.432716

**Authors:** Paulino Ramirez, Wenyan Sun, Gabrielle Zuniga, Elizabeth Ochoa Thomas, Sarah L. DeVos, Bradley Hyman, Gabriel Chiu, Ethan R. Roy, Wei Cao, Miranda Orr, Virginie Buggia-Prevot, William J. Ray, Bess Frost

**Affiliations:** Barshop Institute for Longevity and Aging Studies, San Antonio, Texas; Glenn Biggs Institute for Alzheimer’s and Neurodegenerative Diseases, San Antonio, Texas; Department of Cell Systems and Anatomy, University of Texas Health San Antonio, San Antonio, Texas; Department of Neurology, MassGeneral Institute for Neurodegenerative Disease, Massachusetts General Hospital, Harvard Medical School, Charlestown, Massachusetts; Huffington Center on Aging, Baylor College of Medicine, Houston, TX; Department of Internal Medicine, Wake Forest School of Medicine, Winston-Salem, NC; WG Hefner VA Medical Center, Salisbury, NC; The Neurodegeneration Consortium, Therapeutics Discovery Division, University of Texas MD Anderson Cancer Center, Houston, TX

**Keywords:** Alzheimer’s disease, tauopathy, aging, tau, retrotransposon, endogenous retrovirus, neurodegeneration, mice

## Abstract

Transposable elements comprise almost half of the mammalian genome. A growing body of evidence suggests that transposable element dysregulation accompanies brain aging and neurodegenerative disorders, and that transposable element activation is neurotoxic. Recent studies have identified links between pathogenic forms of tau, a protein that accumulates in Alzheimer disease and related tauopathies, and transposable element-induced neurotoxicity. Starting with transcriptomic analyses, we find that age- and tau-induced transposable element activation occurs in the mouse brain. Among transposable elements that are activated at the RNA level in the context of brain aging and tauopathy, we find that the endogenous retrovirus (ERV) class of retrotransposons is particularly enriched. We show that protein encoded by *Intracisternal A-particle*, a highly active mouse ERV, is elevated in brains of tau transgenic mice. We further demonstrate that brains of tau transgenic mice contain increased DNA copy number of transposable elements, raising the possibility that these elements actively retrotranspose in the context of tauopathy. Taken together, our study lays the groundwork for future mechanistic studies focused on transposable element regulation in the aging mouse brain and in mouse models of tauopathy, while providing support for therapeutic approaches targeting transposable element activation for Alzheimer disease and related tauopathies.

## INTRODUCTION

Approximately 40% of the human genome is composed of retrotransposons^1^. Long and Short Interspersed Nuclear Elements (LINEs, and SINEs, respectively), and Long-Terminal Repeat (LTR)-containing elements make up the major classes of retrotransposons^2^. Intact LINE elements are 6-8 kb and harbor two open reading frames encoding proteins with RNA binding, nucleic acid chaperone, endonuclease and reverse transcriptase activities^3,4^. LINE-encoded proteins function to reverse transcribe LINE and SINE RNA and subsequently insert the newly generated LINE or SINE DNA copy into genomic DNA. Endogenous retroviruses (ERVs) are structurally similar to exogenous retroviruses and are the predominant class of LTR-containing retrotransposons in mammals. Like exogenous retroviruses, the polyprotein encoded by intact ERVs encodes matrix and capsid proteins, as well as a protease, reverse transcriptase, integrase, and envelope proteins^5,6^. Reverse transcription of the ERV RNA is thought to occur inside the capsid, after which the integrase facilitates reinsertion of the new ERV DNA copy into genomic DNA^7^. A simplified overview of the lifecycle of an intact, autonomous LINE or LTR retrotransposon is depicted in **Figure 1a**.

**Figure 1.**
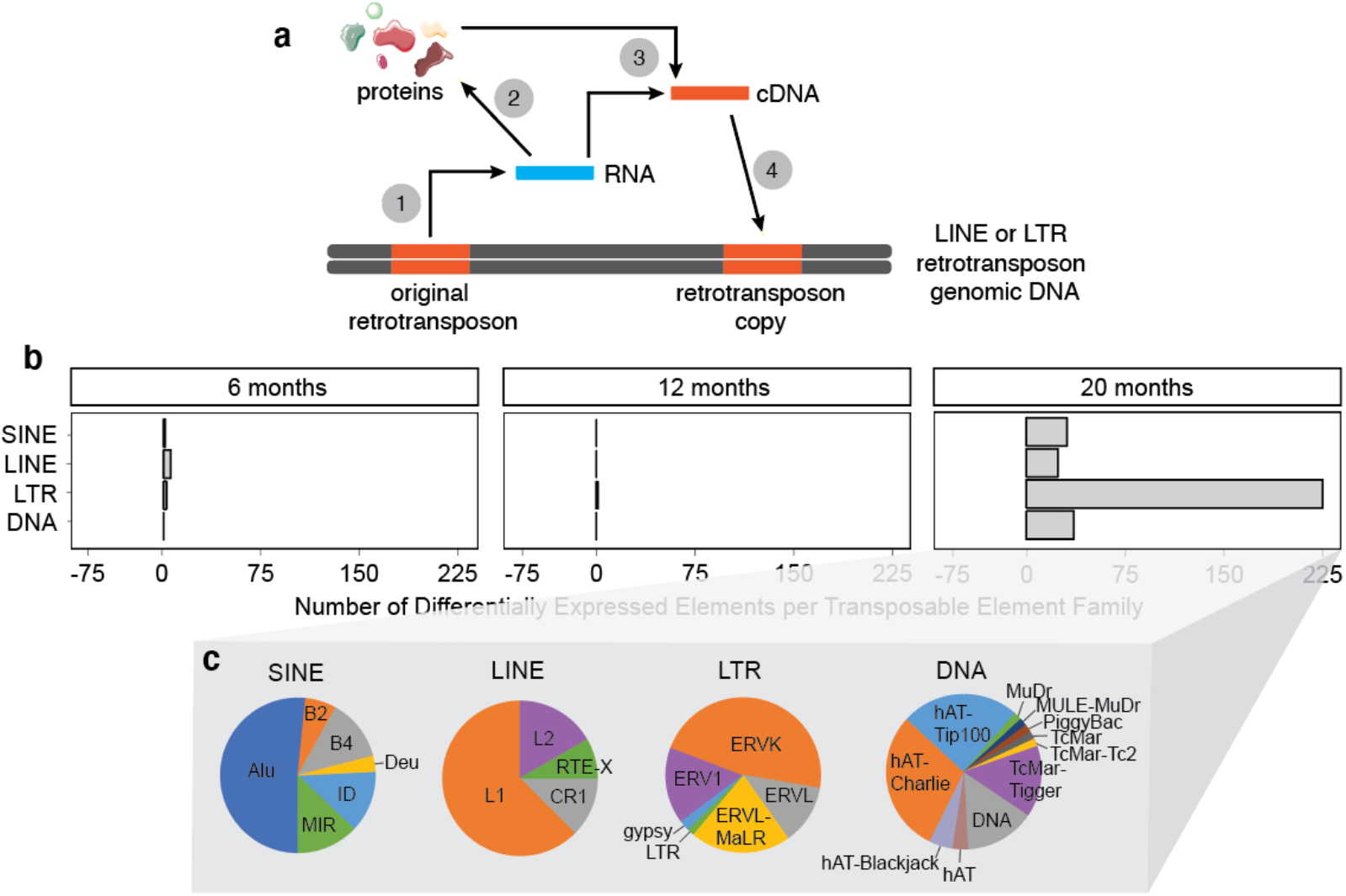
Age-dependent increase in transposable element RNA levels in the mouse forebrain. **a.** Lifecycle of an intact, autonomous retrotransposon: Retrotransposon DNA is transcribed into RNA (1), which encodes proteins that are needed for reverse transcription and subsequent integration into genomic DNA (2). Using retrotransposon-encoded proteins, RNA is reverse transcribed into cDNA that can either exist in an episomal state (3) or can be inserted into genomic DNA (4). **b.** The number of differentially expressed transposable elements in the mouse forebrain from six to twenty months old based on RNA-seq. Differentially expressed transposable elements were grouped into families and represented on the graph. **c.** The proportion of different types of transposable element subfamilies within each transposable element family. n=8-11 mice per age. An adjusted *P* value of <0.05 was considered significant.

It is now understood that transposition can provide an evolutionary advantage by driving genome expansion and evolutionary diversity; however, unchecked transposition can cause DNA breaks and genomic instability, genetic polymorphisms, and insertional mutations that can be detrimental to the organism^8^. While the human and mouse genomes share a similar density of transposable elements, mice have retained a high level of transposon activity over the course of evolution, while most transposable elements in humans have become transpositionally inert^1^, with the exception of ~35-40 subfamilies of L1, Alu, and SVA^9^. In addition to mutations induced by transposition, proteins, single- and double-stranded RNAs, and episomal DNA produced from retrotransposons can also impact cellular function^10^ (**Fig. 1a**). When considering potential toxicity of retrotransposon activation in a given system, retrotransposon-derived products must thus be considered in addition to consequences of retrotransposition to genomic DNA.

The “transposon theory of aging” posits that the cellular control systems that keep transposable elements in check decline with age, and that loss of transposable element control contributes to age-related decline in tissue function^11–14^. Retrotransposon activation has been documented in the context of physiological aging of the *Drosophila* fat body and brain^13,14^. In the liver and muscle of aging mice, several retrotransposons are elevated at both the RNA and DNA levels, suggesting that age-associated transposition also occurs in somatic cells of mammals^12^. We and others have recently reported that retrotransposons become activated in *Drosophila* models of tauopathy and in post-mortem brains of patients with Alzheimer’s disease or progressive supranuclear palsy, a “primary” tauopathy^15,16^. Alzheimer’s disease and related tauopathies are aging-associated human neurodegenerative disorders that involve the deposition of pathological forms of tau protein^17^. Studies in fly models of tauopathy suggest that retrotransposon activation is a consequence of tau-induced breakdown of heterochromatin- and piRNA-mediated retrotransposon silencing, is a causal driver of neurodegeneration, and can be suppressed by antiretroviral therapy^15^.

In the current study, we report that aging- and tau-induced activation of retrotransposons, particularly ERVs, occurs in the mouse brain. Our age-dependent analyses of retrotransposon activation at the RNA, protein, and DNA levels provides a foundation for future mechanistic studies, and support ongoing clinical trials and drug development focused on blocking retrotransposon activation in Alzheimer’s disease and related tauopathies.

## RESULTS

### Age-dependent increase of transposable element transcripts in the mouse brain

To determine if transposable elements are differentially expressed as a consequence of aging in the adult mouse brain, we analyzed transposable element transcript levels in forebrain lysates from mice aged to six, twelve and twenty months of age based on RNA sequencing (RNA-seq). Among the transposable elements that are differentially expressed in the mouse brain by twenty months, we observe that 92.7% increase with age (**Fig. 1b**, **Supplemental Dataset 1**). Among LINE, LTR, and DNA families of transposable elements that are increased in brains of twenty-month-old mice, we find that the LTR family is particularly affected and that ERVs are the predominant family of LTR retrotransposon that increases with age (**Fig. 1c**).

### Age-dependent increase of transposable element transcripts in brains of tau transgenic mice

Based on previous studies reporting elevated transcript levels of select retrotransposons in human tauopathies as well as *Drosophila* models of tauopathy^15,16^, we sought to determine if retrotransposon transcripts are elevated in mouse models of tauopathy. We analyzed RNA-seq datasets across several disease stages from three different mouse models of tauopathy: rTg4510^18^, JNPL3^19^, and PS19^20^.

rTg4510 mice have a *Camk2a*-driven 13-fold overexpression of a familial tauopathy-associated mutant form of human tau, *tau^P301L^*, on a mixed 129S6, FVB genetic background. Tau tangles can be detected by four months of age, while neuronal and synaptic loss occur by six and nine months, respectively^21,22^ (**Fig. 2a**). As females exhibit a higher degree of tau pathology and behavioral deficits than males in this model^23^, we analyzed transposable element transcript levels in cortical lysates from female control and rTg4510 mice at three, six, and nine months of age. We detect a general increase in transposable element transcripts by nine months of age in rTg4510 tau transgenic mice (**Fig. 2b**, **Supplemental Fig. 1a**, **Supplemental Dataset 1**). Similar to our findings in the context of physiological mouse brain aging, LTR retrotransposons have the largest increase in rTg4510 mice, with ERVs as the predominant LTR that is elevated (**Fig. 2c**).

**Figure 2.**
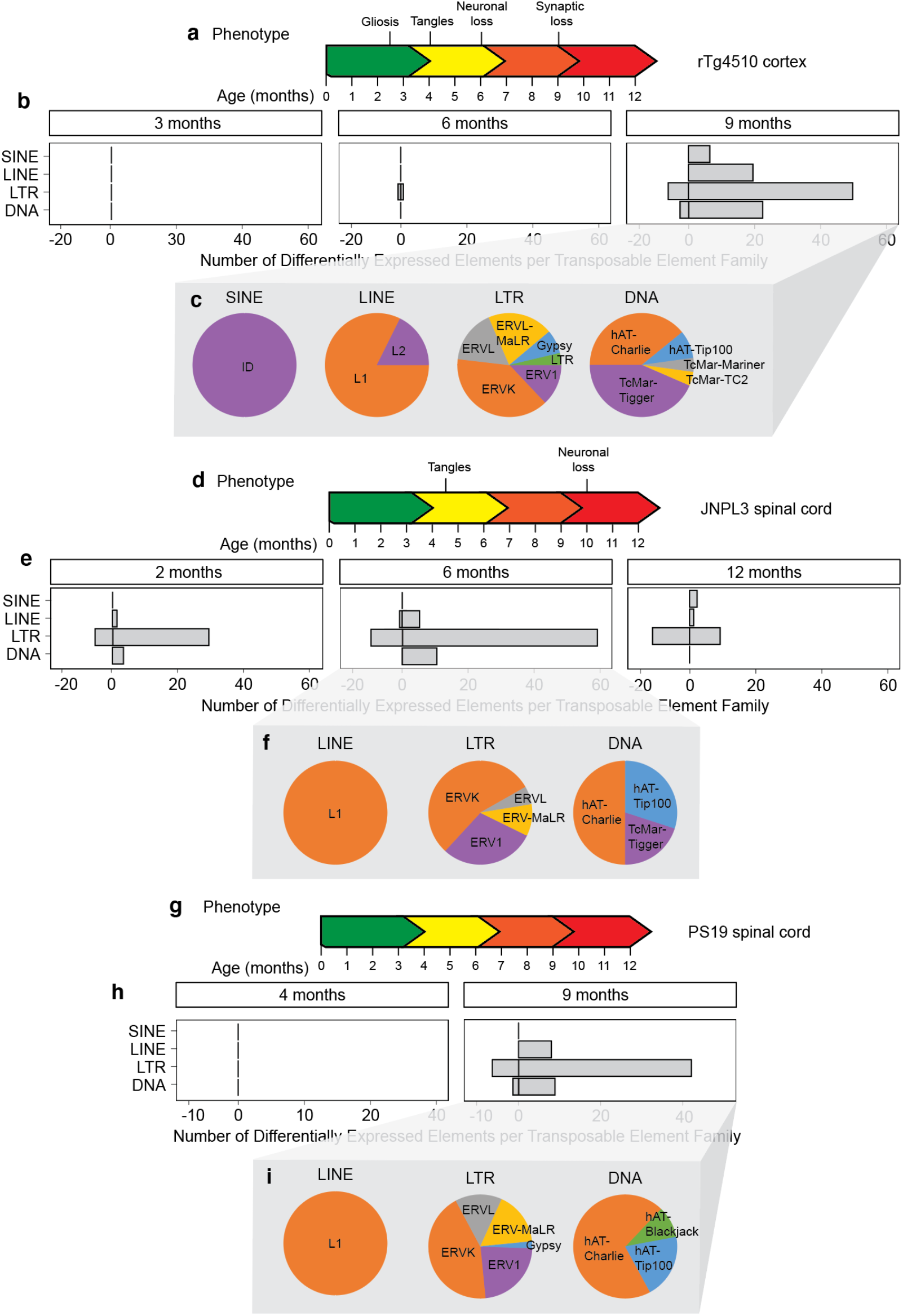
Age-dependent analysis of transposable element RNA levels in three different mouse models of tauopathy. **a.** The rTg4510 mouse model of tauopathy. **b.** Differentially expressed transposable elements in the female rTg4510 mouse cortex from three to nine months based on RNA-seq. Differentially expressed transposable elements were grouped into families and represented on the graph. n=5-6 biological replicates per age. **c.** The proportion of different types of differentially expressed transposable element subfamilies within each transposable element family. **d.** The JNPL3 mouse model of tauopathy. **e.** Differentially expressed transposable elements in the female JNPL3 spinal cord from two to twelve months based on RNA-seq. n=3-6 biological replicates per age. **f.** The proportion of differentially expressed transposable element subfamilies within each transposable element family. **g.** The PS19 mouse model of tauopathy. **h.** Differentially expressed transposable elements in the male PS19 spinal cord at four and nine months based on RNA-seq. Differentially expressed transposable elements were grouped into families and represented on the graph. n=5-7 biological replicates per age. **I.** The proportion of differentially expressed transposable element subfamilies within each transposable element family. An adjusted *P* value of <0.05 was considered significant.

The JNPL3 model features mouse prion promoter-driven expression of human tau^P301L^ on a C57Bl/6, DBA/2, SW mixed genetic background. JNPL3 mice produce human tau protein at a level similar to endogenous mouse tau. As neuronal loss in the spinal cord is a predominant feature of this model^19^, we utilized publicly available RNA-seq data from spinal cord of homozygous JNPL3 mice at two, six, and twelve months of age. RNA-seq data was only available for females, as males exhibit a delayed and variable phenotype in this model. Tau tangles can be detected by six to seven months in the brainstem and spinal cord in female homozygotes, while loss of motor neurons occurs at approximately ten months^19^ (**Fig. 2d**). We detect an overall elevation of LTR elements as early as two months in this model, which precedes tau tangle formation. The increase in retrotransposon transcripts is elevated further at six months, after which we detect a moderate decline by twelve months (**Fig. 2e**, **Supplemental Fig. 1b**, **Supplemental Dataset 1**). Again, we find ERVs to be the predominant class of LTR retrotransposons that are elevated in this model of tauopathy (**Fig. 3f**).

**Figure 3.**
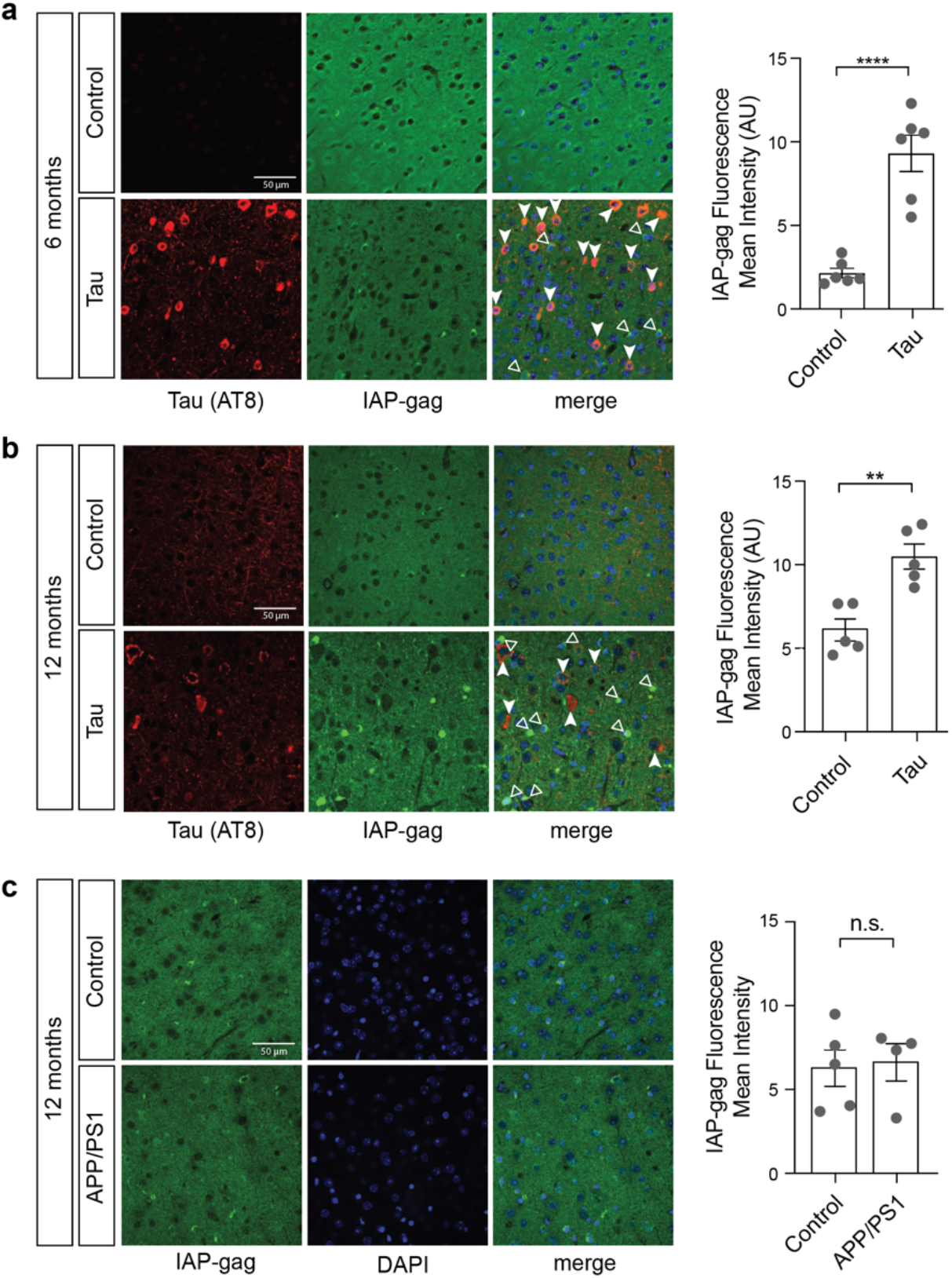
*IAP*-encoded gag protein in rTg4510 and APP/PS1 mouse models. Frontal cortices of rTg4510 mice at six (**a**) and twelve (**b**) months were stained with antibodies recognizing IAP-encoded gag and tau AT8. **c.** Frontal cortices of APP/PS1 mice were stained with antibodies recognizing IAP-encoded gag. n=4-6 biological replicates per genotype, per age. Filled-in arrows indicate tau AT8-positive cells, while empty arrows indicate IAP-gag-positive cells. **p<0.01, ****p<0.0001, unpaired t-test. Error bars=SEM.

The PS19 model features a five-fold overexpression of the familial tauopathy-associated tau^P301S^ mutation driven by the mouse prion promoter on a C57Bl/6, C3H genetic background. The original publication reports presence of neurofibrillary tangles in the brain and spinal cord by six months and neuronal loss by nine months^20^. As reported by others^24,25^, our colony has delayed pathology compared to the original line, with higher levels of pathology in the spinal cord compared to the hippocampus. We analyzed RNA-seq data from spinal cord of male PS19 mice at four and nine months of age, as males generally have more consistent presentation of tau pathology in this model (**Fig. 2g**). While we do not detect any transposable elements that are differentially expressed at four months of age, we observe an increase in retrotransposon transcripts in the spinal cord of PS19 mice by nine months (**Fig. 2h**, **Supplemental Fig. 1c**, **Supplemental Dataset 1**). Similar to our findings in human tauopathy^15^, physiological mouse brain aging, and rTg4510 and JNPL3 mice, we find that LTR retrotransposons are the predominant class of transposable elements that are elevated at the transcript level in PS19 spinal cord, and that ERVs are the predominant type of LTR retrotransposon that is elevated. Collectively, these data demonstrate a general age-dependent increase in retrotransposon transcript levels in tau transgenic mice despite different promoters, tissues, sexes, genetic backgrounds, and transgenic tau dosage, demonstrating that tau-induced ERV activation at the RNA level is robustly conserved among various mouse models.

### Increase in protein encoded by an active ERV in brains of tau transgenic mice

As RNA-seq revealed an age-dependent increase in ERV retrotransposons in the context of normal aging and three different mouse models of tauopathy, we next asked if ERV-encoded proteins can be detected in the brain of tau transgenic mice. We focused on the “<u>g</u>roup <u>a</u>nti<u>g</u>ens” (gag) polyprotein encoded by a highly active mouse ERV, *Intracisternal A Protein* (*IAP*), which has approximately 2,800 complete or near-complete copies in the reference C57BL/6 genome^26^. As we do not detect a significant increase in RNA levels of ERVs prior to nine months of age in rTg4510 mice based on RNA-seq, we analyzed *IAP*-encoded protein at six months (prior to elevation in ERV RNA) and twelve months (after elevation in ERV RNA) using an antibody^27^ that detects IAP-gag. We find that IAP-gag is significantly elevated in frontal cortex of rTg4510 mice at both six and twelve months of age (**Fig. 3a, b**). Interestingly, the cells harboring elevated gag protein levels are not the cells harboring a disease-associated form of human tau that is phosphorylated at serine 202 and threonine 205 (AT8), suggesting that tau-induced ERV activation is non-cell autonomous.

To determine if retrotransposon activation is a common feature of all mouse models of brain protein aggregation, we visualized IAP-gag protein in brains of a mouse model carrying human transgenes for mutant *amyloid precursor protein* and *presenilin 1* (APP/PS1). Cortical deposition of amyloid plaques can be detected by six weeks in this model^28^. We do not detect differences in *IAP*-encoded gag at twelve months of age in the frontal cortex of APP/PS1 mice (**Fig. 3c**), suggesting that retrotransposon activation does not occur in all mouse models of protein aggregation.

### The IAP-encoded gag polyprotein is processed to a mature form in brains of tau transgenic mice

Intact copies of *IAP* encode a gag-pro-pol polyprotein precursor and env, which facilitates budding of the endogenous viral particle from the plasma membrane. Many copies of *IAP* lack a functional *env* domain, and bud into the cisternae of the endoplasmic reticulum rather than into the extracellular space^29^. The *IAP*-encoded gag-pro-pol polyprotein undergoes a series of proteolytic cleavages by the pro-encoded protease. Gag is further processed to generate capsid, matrix, and nucleocapsid proteins^30^.

We took a biochemical approach to determine if the *IAP*-encoded polyprotein is processed to a mature form in brains of rTg4510 mice. As we detected elevated levels of *IAP*-encoded gag in rTg4510 mice by six months of age, we extended our earlier timepoint to two months of age. We generated an antibody that recognizes the polypeptide sequence corresponding to the capsid protein cleaved from the gag precursor. While this epitope is present in multiple cleavage products of the gag-pro-pol precursor, and will thus detect multiple stages of processing, proteins generated by cleavage of the precursor can be differentiated by size (**Fig. 4a**). While *IAP*-encoded polyprotein processing at two months does not differ between control and tau transgenic mice (**Fig. 4b**), twelve-month-old rTg4510 mice have a significant decrease in a ~120 kD band, with a concomitant increase in bands at ~52 kD and ~24 kD, consistent with protease-mediated cleavage of an *IAP*-encoded polyprotein and generation of a mature gag capsid protein (**Fig. 4c, d**). These data suggest that the polyprotein encoded by *IAP* is processed to a greater extent in the rTg4510 model, consistent with activation.

**Figure 4.**
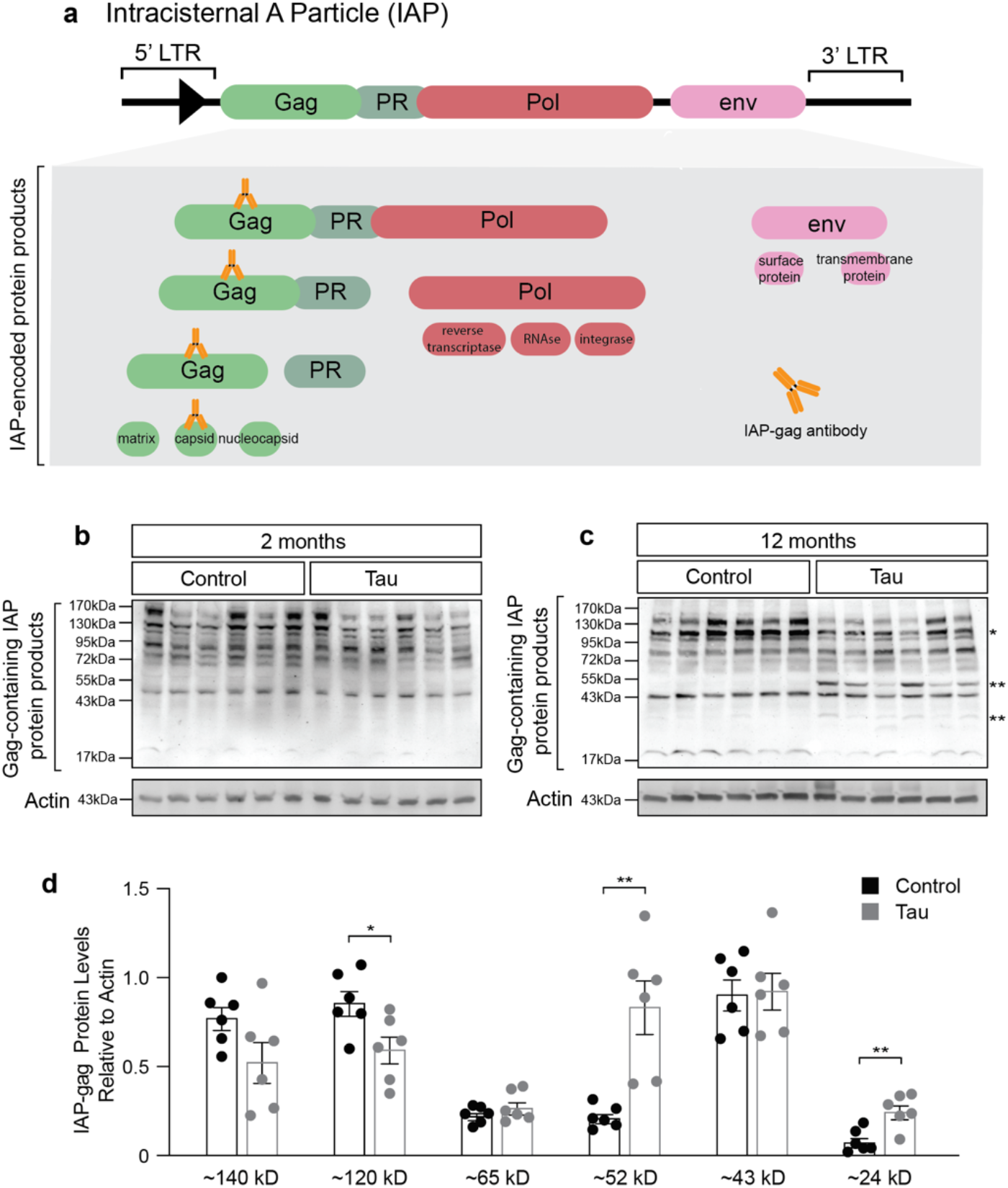
IAP polyprotein processing is increased in brains of rTg5410 mice at twelve months of age. **a.** Intact copies of the *IAP* gene encode a gag-pro-pol precursor and env. The gag-pro-pol precursor is processed into multiple protein products. The antibody used for western blotting recognizes the capsid protein that is produced from gag. IAP polyprotein processing based on immunoblotting with an antibody recognizing the gag-encoded capsid protein in lysates from rTg4510 mouse brain compared to control at two (**b**) and twelve (**c**) months of age. **d.** At twelve months of age, rTg4510 mice have significantly decreased levels of a band corresponding to a ~120 kD gag-pro-pol precursor, and significantly increased levels of ~52 kD and ~24 kD gag-containing proteins. n=6 biological replicates, two-sided t-tests shown on the same graph for visual ease. *p<0.05, **p<0.01. Error bars=SEM.

### Transposable element DNA copy number is elevated in tau transgenic mice

As retrotransposition occurs through a copy-and-paste mechanism, each mobilization event increases the total number of DNA copies of a given element by one. Having identified retrotransposons that are elevated at the RNA and protein levels in brains of tau transgenic mice, we next asked if retrotransposon DNA copy number is elevated in brains of rTg4510 mice. We quantified DNA copy number of retrotransposons in cortical brain tissue from control and rTg4510 mice using a custom NanoString codeset. The codeset consists of probes that detect DNA of both autonomous and non-autonomous L1, L2, ERV1, ERVK, ERVL, ERVL-MaLR and solo LTRs. As subfamily members within a transposable element family share a high degree of sequence similarity, many of the probesets recognize multiple subfamily members based on BLAST analysis (**Supplemental Dataset 2**). For each probeset, DNA copy number was normalized to ten different invariant regions of the mouse genome (**Supplemental Dataset 2**).

At two months of age, rTg4510 have a significant increase in L1 probeset A compared to controls (**Fig. 5a**, **Supplemental Dataset 2**). At twelve months of age, we detect a three-fold elevation in the elements recognized by L1 probeset A, as well as significant increases in ERVK probesets A-E and ERV1 probeset A in brains of rTg4510 mice compared to controls (**Fig. 5b, c**, **Supplemental Dataset 2**). Retrotransposon DNA copy number does not significantly differ between two and twelve months in control mice (**Supplemental Dataset 2**). Taken together, these data indicate that a subset of retrotransposon RNAs are actively reverse transcribed into DNA in brains of rTg4510 transgenic mice, and that tau-induced increase in retrotransposon DNA copy number is age-dependent.

**Figure 5.**
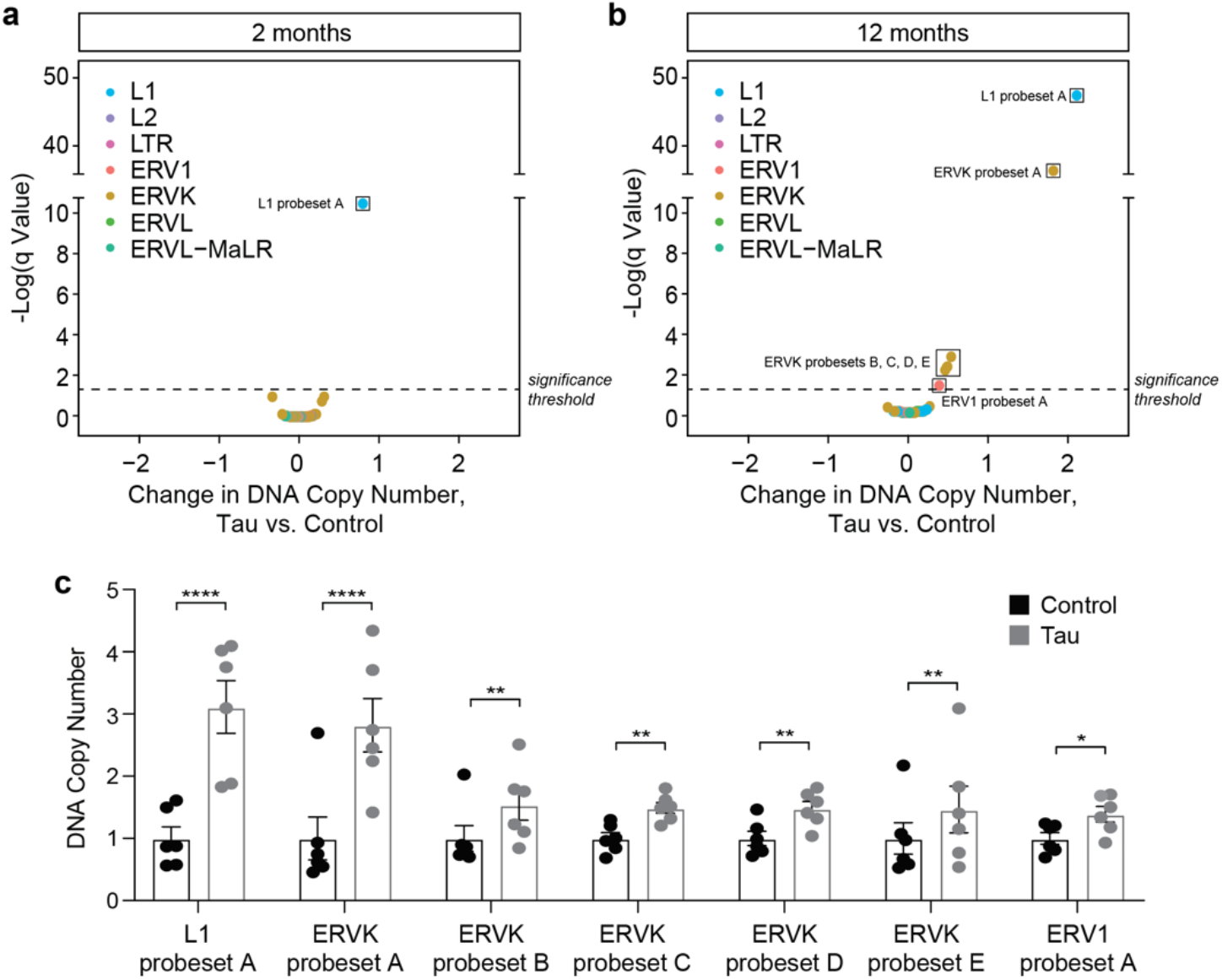
NanoString-based transposable element DNA copy number analysis in the cortex of rTg4510 mice. DNA copy numbers of transposable elements in cortex of rTg4510 vs. control mouse brain at two (**a**) and twelve (**b**) months of age. Elements that are significantly different between rTg4510 and control are boxed. False-discovery rate, q=1%. **c.** Individual data points for elements that are significantly increased in rTg4510 mouse brain at twelve months of age compared to control. n=6 females per genotype, per age. Error bars=SEM. Multiple *t*-tests, *q<0.05, **q<0.01, ****q<0.000001.

## DISCUSSION

Multiple previous reports suggest that retrotransposons are activated in the brain as a consequence of physiological aging in *Drosophila*^13,14^, and that pathogenic forms of tau also activate retrotransposons in *Drosophila* models of tauopathy and in postmortem human brain tissue from patients with tauopathies, including Alzheimer’s disease^15,16^. In the current study, we investigate the extent of retrotransposon activation from young to old age in the mouse brain and over the course of disease in three different mouse models of tauopathy.

We find that the directionality of differentially expressed retrotransposons is predominantly increased in the context of both aging and tauopathy in the mouse brain. This is in slight contrast to previous studies that identify retrotransposons that are both increased and decreased with aging and tauopathy in the *Drosophila* brain, as well as in postmortem human brains of patients with late-stage Alzheimer’s disease or progressive supranuclear palsy^15^. Similar to human Alzheimer’s disease and tau transgenic *Drosophila*, we find an overrepresentation of ERVs among the classes of transposable elements that are elevated at the transcript level in the context of aging and tauopathy in the mouse brain. Increased transcript levels of ERVs are also present in human disorders including but not limited to amyotrophic lateral sclerosis^31,32^, multiple sclerosis^33^, and various types of cancers^34^.

In addition to identifying retrotransposons that are elevated at the transcript level in the context of tauopathy, we find that the gag capsid protein encoded by *IAP*, a highly active mouse Class II ERV, is elevated in the brain of rTg4510 tau transgenic mice. While it is possible that differential IAP processing precedes differential expression of *IAP*, we speculate that our ability to detect elevated levels of IAP-encoded protein prior to our ability to detect elevated IAP transcript levels in brains of rTg4510 mice is a consequence of sequencing depth. While we do not currently know the consequences of elevated levels or differential processing of IAP in brains of tau transgenic mice, a previous study has found that transgenic expression of a human ERV-encoded envelope protein is sufficient to induce motor neuron degeneration in mice^35^.

Having established that ERVs are elevated at the RNA and protein levels in brains of tau transgenic mice, we next asked if brains of tau transgenic mice have increased retrotransposon DNA content. We find that subfamily members of LINE and ERV families are significantly increased in brains of rTg4510 tau transgenic mice compared to control. As NanoString-based DNA copy number variation analysis does not discriminate between genomic and episomal DNA, we do not currently know whether the extra retrotransposon DNA copies are integrated into the genome and/or exist in an episomal state. Determining the proportion of extra retrotransposon copies that are genomic versus episomal is an important next step, as genomic insertions generate novel mutations, while episomal DNA could drive a viral response as described in the context of aging, senescence and activation of LINE-1 elements in somatic tissues^36^.

Cells have multiple mechanisms to safeguard the genome against transposable element activation. Transposable element transcription is limited by enrichment of heterochromatin around retrotransposon DNA, while retrotransposon transcripts are degraded by small RNAs termed piwi-interacting RNAs (piRNAs) and endogenous small interfering RNAs (esiRNAs)^37^. We have previously reported that pathogenic forms of tau disrupt heterochromatin- and piRNA-mediated silencing of retrotransposons in the *Drosophila* brain^15^. While our current study does not address mechanistic links between pathogenic forms of tau and retrotransposon activation in the mouse brain, depletion of heterochromatin protein 1 in motor neurons of the spinal cord in the JNPL3 mouse model of tauopathy^38^ is consistent with our overall hypothesis that pathogenic tau-induced heterochromatin decondensation drives retrotransposon activation. In addition, studies in tau knockout mice suggest that maintaining the integrity of pericentromeric heterochromatin is a physiological function of tau^39^. Investigation into heterochromatin- and piRNA-mediated control over retrotransposons in the aging mouse brain and in mouse models of tauopathies will be the subject of future studies.

It has been 17 years since the last drug was approved for the symptomatic treatment of Alzheimer’s disease. Treatment pipelines for primary tauopathies are in their infancy, with only a few having disease-modifying potential. Pathogenic forms of tau and downstream cellular mediators of neuronal death represent potential therapeutic targets that have been largely unexploited. Due to the similarities between retroviruses and retrotransposons, antiretroviral drugs that interfere with HIV-1 replication have recently gained momentum as therapeutic approaches in neurodegenerative disorders, with promising results from a Phase 2 clinical trial for ALS (NCT02868580)^40^ and recent initiation of Phase 1 (NCT04500847) and Phase 2 trials for Alzheimer’s disease (NCT04552795). Taken together, our data confirms that aging- and tau-induced transposable element activation is conserved in the mouse central nervous system, and further strengthens the case for transposable element activity as a therapeutic target for human tauopathies.

## METHODS

### Mouse Models

#### rTg4510

The founder rTg4510 and non-transgenic control mice were obtained from the Jackson Laboratory (IMSR_JAX:024864). Mice were group-housed in a pathogen-free mouse facility with *ad libitum* access to food and water on a twelve-hour light/dark cycle. After anesthesia with 2% isoflurane, cardiac perfusion was performed using 2X PhosSTOP phosphatase inhibitors (Roche, Indianapolis, IN, USA) and 1X complete protease inhibitors (Roche) in PBS (Thermo Fisher Scientific, Waltham, MA, USA). All experimental procedures were performed according to the NIH Guidelines for the Care and Use of Laboratory Animals and were approved by the Institutional Animal Care and Use Committee of the University of Texas MD Anderson Cancer Center.

#### PS19

The founder PS19 mice were obtained from the Jackson Laboratory (IMSR_JAX:008169). Heterozygotic offspring were bred and maintained at Baylor College of Medicine. Wild-type littermates were used as controls. Mice were housed 4-5 per cage in a pathogen-free mouse facility with *ad libitum* access to food and water on a twelve-hour light/dark cycle. Male mice were analyzed in this study. All procedures were performed in accordance with NIH guidelines and approval of the Baylor College of Medicine Institutional Care and Use Committee.

### RNA Sequencing and Analysis

RNA was isolated from hippocampi of rTg4510 and control, and spinal cord of PS19 and control mice using the Qiagen RNeasy Mini kit and Qiagen TissueRuptor (Qiagen, Germantown, MD, USA) based on the manufacturer’s recommendations. RNA from n=17-18 rTg4510 female mice and controls per timepoint was sequenced using an Illumina HiSeq 4000 with 76 bp paired-end reads at ~30 million reads per sample. RNA from 5-7 PS19 male mice was sequenced using an Illumina HiSeq 6000 with 150 bp paired-end reads at ~20 million reads per sample. For transposable element analysis in the aging mouse brain RNA-seq data from control mice (B6C3HF1) were downloaded from AMP-AD dataset syn3157182 (n=1 male, 7 females at 6 months, n=4 males and 4 females at 12 months, and n=6 males and 5 females at 20 months). RNA-seq data from spinal cord of n=3-6 JNPL3 and control female mice per timepoint were downloaded from AMP-AD dataset syn3157182. B6C3HF1, JNPL3 and control mice were sequenced on an Illumina HiSeq 2000 with 101 bp paired end reads at ~50 million reads per sample. RNA libraries were prepared with Illumina TruSeq library kits. Each tau model has its own set of age-matched control samples.

FastQ files for each experiment were processed with Trimmomatic^41^ v.0.36 (SLIDINGWINDOW:5:20 MINLEN:50) to remove low quality reads and trim adapter sequences. FastQC^42^ was then used to obtain pre and post quality control processing statistics for the FastQ files. Trimmed paired reads were aligned with STAR^43^ v.2.7.1a to the mm10 mouse genome using parameters suggested by TEtranscripts^44^ (--outFilterMultimapNmax 100, --winAnchorMultimapNmax 100). Transposon counts for each sample were then obtained using TEtranscripts v.2.0.3 (default parameters) and its supplied mouse mm10 genome transposon annotation file.

Differential transposon expression analysis was completed with DESeq2^45^. Pairwise comparisons were performed with the DESeq2 Wald test on rTg4510, JNPL3 and PS19 and their age- and model-matched controls. To quantify transposable element RNA changes as a consequence of aging, differential transposon expression was performed as a time series analysis (test = LRT, full model = Sex + Age, reduced model = ~1) at six, twelve and twenty months. Differentially expressed transposons from the LRT test were clustered into similar expression patterns across the different age groups using DEGreports. The three clusters correspond to the age at which expression of the transposable element was at its highest. Genes with an adjusted *P* value of <0.05 were considered significantly differentially expressed^32^.

Barplots and heatmaps were generated using the R packages ggplot2^46^ and pHeatmap^47^, respectively. Transposable elements were grouped into families and subfamilies using the TEtranscripts^44^ supplied annotation information.

### Development of the IAP-gag Antibody

The nucleocapsid sequence corresponding to the capsid protein that is processed from the IAP gag precursor was sent to GenScript for antibody development:

LTGQGAYSADQTNYHWGAYAQISSTAIRRWKGLSRAGETTGQLTKVVQGPQESFSDFVARMTEAAERIFGESEQAAPLIEQLIYEQATKECRAAIAPRKNKGLQDWLRVCRELGGPLTNAGL

The peptide was expressed in *E. coli* and used to immunize two rabbits for antibody production. The GeneScript proprietary OptimumAntigenTM design tool was used to measure the antigen against several protein databases to enhance the desired antibody and epitope specificity. GenScript utilized codon optimization and BacPowerTM bacterial protein expression technology and proprietary adjuvant to increase the efficiency of protein expression and animal immunization. An ELISA titer of over 1:128,000 and target protein binding were validated by immunoprecipitation and western blotting using the positive control with protein immunogen by GenScript. The final product was 0.5 ml of pre-immune serum at 1.5-6 mg/rabbit and 1 mg of the requested peptide.

### Western Blotting

10-20 mgs of frozen mouse brain cortex were homogenized in 60 μl Laemmli sample buffer (Invitrogen). Total protein content was quantified with a BCA protein assay kit (Pierce) and normalized across samples. Lysates containing approximately 40 μg of protein were boiled for 10 min prior to separation by 4-20% SDS-PAGE (BIO-RAD). After transferring at 90V to a nitrocellulose membrane for 90 min at room temperature, Ponceau S staining was performed to ensure that protein loading was equal among samples prior to membrane blocking with 2% milk in PBS with 0.05% Tween (PBSTw) at room temperature for 30 min. The membrane was then incubated with the anti-IAP-gag antibody at 1:500 (GeneScript) overnight at 4°C. Membranes were washed three times with PBSTw and incubated with an HRP-conjugated secondary antibody for 2 h at room temperature. Blots were developed with an enhanced chemiluminescent substrate (Thermo Scientific). After imaging, membranes were stripped and incubated with an anti-Actin antibody at 1:1,000 dilution (#JLA20, DSHB) overnight at 4°C. The following day, membranes were washed and incubated with an HRP-conjugated secondary antibody, and imaged with an enhanced chemiluminescent substrate. Band densitometry was performed using ImageJ.

### Immunostaining

40 μm mouse brain sections were stored at 20°C in cryoprotectant prior to use. Individual sections were transferred to PBS in single wells of a 24 well plate. After washing three times with PBS, sections were incubated overnight with the IAP-gag primary antibody^27^ (1:100, a gift from Dr. Brian Cullen) and AT8 (1:200, #MN1020, Fisher) in PBS + 0.3% Triton + 2% BSA at 4°C overnight. After three washes in PBS+Triton, sections were incubated with Alexa Fluor 488- or 555-conjugated IgG (1:200, Invitrogen) at room temperature for 2 h. After washing, the sections were mounted onto microscope slides and then coverslipped with Fluoromount G with DAPI. Immunofluorescence was visualized on a Zeiss confocal microscope and analyzed with ImageJ. Brain sections from twelve month old rTg4510 and APP/PS1 models used for IAP-gag immunostaining were generated from mice used in a previous study^48^.

### NanoString

In collaboration with NanoString Technologies, Inc., we created a custom nCounter codeset consisting of 188 probes targeting 89 mouse transposable elements and 10 internal controls. Two probes were designed for each transposable element region and one probe was designed for each internal control (see **Supplemental Dataset 2** for codeset sequences and data). Single copy or low copy genes were used as internal controls. To identify transposable element family members that are recognized by each probe, probe sequences were BLASTed^49^ to a database consisting of mouse transposable elements from Repbase^50^ and transposable elements present in the TEtranscripts^44^ mouse transposable element GTF file. Family members with >85% identity to the probe sequence and >75 bp length were considered hits. 3-5 μg of DNA was isolated from frozen cortex of 2- and 12-month-old mice according to the manufacturer’s protocol (DNeasy Blood & Tissue Kit, Qiagen, Cat No. 69504) and diluted with 10 mM Tris. Prior to hybridization, diluted DNA was processed in a Covaris Focused-ultrasonicator to produce 200 bp fragments. DNA fragmentation was confirmed using an Agilent 2100 Bioanalyzer. 300 ng of fragmented DNA was isolated via ethanol precipitation and used for NanoString nCounter Copy Number Assays. nCounter SPRINT profiler was used to quantify copy number of each target. Raw retrotransposon probe counts were normalized to ten probes recognizing invariant regions of the mouse genome (**Supplemental Table 2**) using nSolver 4 in order to correct for technical variability. Transposable elements with significantly different DNA copy numbers between rTg4510 and control were detected by multiple *t*-tests assuming equal standard deviation between samples, followed by the two-stage linear step-up procedure of Benjamini, Krieger and Yekutieli with a q value = 0.05 using GraphPad Prism v8.4.2^51^ for MacOS (GraphPad Software, San Diego, CA USA). Volcano plots for DNA copy change were made using ggplot2^52^.

## SUPPLEMENTAL DATASETS

**Supplementary Dataset 1 | Differential expression analysis in the aging mouse brain and in rTg4510, JNPL3, and PS19 tau transgenic mice.** Transposable elements that are significantly differentially expressed based on an adjusted *P* value of <0.05 are highlighted in yellow. For the aging mouse brain, a positive log2 fold change indicates increased transcript levels as a consequence of aging. For mouse models of tauopathy, a positive log2 fold change indicates increased transcript levels in the tauopathy model compared to its age-matched control.

**Supplementary Dataset 2 | NanoString codeset sequences and raw data.** Each probeset consists of two probes designed to detect members of the indicated subfamily. Retrotransposon probe sequences and invariant control probe sequences are listed in the “**Probe Sequence**” sheet. The “**Probes BLAST**” sheet includes the list of retrotransposon family members recognized by each probeset. Retrotransposon DNA copy number for individual mice normalized to internal control genes are included in “**2mo Copy Number, normalized**” and “**12mo Copy Number, normalized**” sheets. Probesets that are significantly different between control and rTg4510 based on a q value of <0.05 are highlighted in yellow in “**2mo rTg4510 vs. Control,**” “**12mo rTg4510 vs. Control**” and “**2mo Control vs. 12mo Control.**”

## FUNDING AND ACKNOWLEDGEMENTS

This study was supported by the Rainwater Foundation (BF, SD, BH), the National Institute of Neurological Disorders and Stroke RF1 NS112391 (BF), the National Institute on Aging RF1 AG057587 (WC), T32 AG00022 (SLD), the National Institute of General Medical Sciences R25 GM095480 (PR) and T32 GM113896 (GZ). Acknowledgement is made to the donors of Alzheimer’s Disease Research, a program of BrightFocus Foundation, for support of this research (WS, SLD). WJR and VBP are supported by the Neurodegeneration Consortium, the Belfer Family foundation and the Oskar Fisher Project. The IAP-gag antibody used for immunofluorescence was a kind gift of Dr. Bryan Cullen.

## CONTRIBUTIONS

PR, WS, and GZ performed experiments, data analysis, and manuscript preparation. EOT and BF participated in data analysis and manuscript preparation. SD and BH generated rTg4510 and APP/PS1 brain tissue for immunostaining. GC, ER, and WC generated RNA-seq data from PS19 mice. VBP and WR generated RNA-seq data from rTg4510 mice and respective controls, and provided brain tissue from rTg4510 and controls for immunostaining, biochemical analysis and NanoString.

## Notes

### Competing Interest Statement

The authors have declared no competing interest.

## REFERENCES

1. Lander, E. S. et al. Initial sequencing and analysis of the human genome. Nature 409, 860–921 (2001).

2. Platt, R. N., Vandewege, M. W. & Ray, D. A. Mammalian transposable elements and their impacts on genome evolution. Chromosome Research (2018) doi:10.1007/s10577-017-9570-z.

3. Martin, S. L. et al. LINE-1 retrotransposition requires the nucleic acid chaperone activity of the ORF1 Protein. J. Mol. Biol. (2005) doi:10.1016/j.jmb.2005.03.003.

4. Feng, Q., Moran, J. V., Kazazian, H. H. & Boeke, J. D. Human L1 retrotransposon encodes a conserved endonuclease required for retrotransposition. Cell (1996) doi:10.1016/S0092-8674(00)81997-2.

5. Hughes, J. F. & Coffin, J. M. A novel endogenous retrovirus-related element in the human genome resembles a DNA transposon: Evidence for an evolutionary link? Genomics (2002) doi:10.1016/S0888-7543(02)96856-4.

6. Nelson, P. N. et al. Demystified … Human endogenous retroviruses. Journal of Clinical Pathology - Molecular Pathology (2003) doi:10.1136/mp.56.1.11.

7. Burns, K. H. Our Conflict with Transposable Elements and Its Implications for Human Disease. Annual Review of Pathology: Mechanisms of Disease (2020) doi:10.1146/annurev-pathmechdis-012419-032633.

8. Hancks, D. C. & Kazazian, H. H. Roles for retrotransposon insertions in human disease. Mobile DNA (2016) doi:10.1186/s13100-016-0065-9.

9. Mills, R. E., Bennett, E. A., Iskow, R. C. & Devine, S. E. Which transposable elements are active in the human genome? Trends in Genetics (2007) doi:10.1016/j.tig.2007.02.006.

10. Beck, C. R., Garcia-Perez, J. L., Badge, R. M. & Moran, J. V. LINE-1 elements in structural variation and disease. Annual Review of Genomics and Human Genetics (2011) doi:10.1146/annurev-genom-082509-141802.

11. Murray, V. Are transposons a cause of ageing? Mutat. Res. DNAging (1990) doi:10.1016/0921-8734(90)90011-F.

12. De Cecco, M. et al. Transposable elements become active and mobile in the genomes of aging mammalian somatic tissues. Aging (Albany. NY). (2013) doi:10.18632/aging.100621.

13. Li, W. et al. Activation of transposable elements during aging and neuronal decline in Drosophila. Nat. Neurosci. (2013) doi:10.1038/nn.3368.

14. Wood, J. G. et al. Chromatin-modifying genetic interventions suppress age-associated transposable element activation and extend life span in Drosophila. Proc. Natl. Acad. Sci. 113, 11277–11282 (2016).

15. Sun, W., Samimi, H., Gamez, M., Zare, H. & Frost, B. Pathogenic tau-induced piRNA depletion promotes neuronal death through transposable element dysregulation in neurodegenerative tauopathies. Nat. Neurosci. 21, 1038–1048 (2018).

16. Guo, C. et al. Tau Activates Transposable Elements in Alzheimer’s Disease. Cell Rep. 5, 2874–2880 (2018).

17. Arendt, T., Stieler, J. T. & Holzer, M. Tau and tauopathies. Brain Res. Bull. 126, 238–292 (2016).

18. Santacruz, K. et al. Medicine: Tau suppression in a neurodegenerative mouse model improves memory function. Science (80-.). (2005) doi:10.1126/science.1113694.

19. Lewis, J. et al. Neurofibrillary tangles, amyotrophy and progressive motor disturbance in mice expressing mutant (P301L)tau protein. Nat. Genet. (2000) doi:10.1038/78078.

20. Yoshiyama, Y. et al. Synapse Loss and Microglial Activation Precede Tangles in a P301S Tauopathy Mouse Model. Neuron (2007) doi:10.1016/j.neuron.2007.01.010.

21. Spires, T. L. et al. Region-specific dissociation of neuronal loss and neurofibrillary pathology in a mouse model of tauopathy. Am. J. Pathol. (2006) doi:10.2353/ajpath.2006.050840.

22. Kopeikina, K. J. et al. Synaptic alterations in the rTg4510 mouse model of tauopathy. J. Comp. Neurol. (2013) doi:10.1002/cne.23234.

23. Yue, M., Hanna, A., Wilson, J., Roder, H. & Janus, C. Sex difference in pathology and memory decline in rTg4510 mouse model of tauopathy. Neurobiol. Aging (2011) doi:10.1016/j.neurobiolaging.2009.04.006.

24. Iba, M. et al. Synthetic tau fibrils mediate transmission of neurofibrillary tangles in a transgenic mouse model of alzheimer’s-like tauopathy. J. Neurosci. (2013) doi:10.1523/JNEUROSCI.2642-12.2013.

25. Zhang, B. et al. The microtubule-stabilizing agent, epothilone D, reduces axonal dysfunction, neurotoxicity, cognitive deficits, and alzheimer-like pathology in an interventional study with aged tau transgenic mice. J. Neurosci. (2012) doi:10.1523/JNEUROSCI.4922-11.2012.

26. Gagnier, L., Belancio, V. P. & Mager, D. L. Mouse germ line mutations due to retrotransposon insertions. Mobile DNA (2019) doi:10.1186/s13100-019-0157-4.

27. Bogerd, H. P., Wiegand, H. L., Doehle, B. P., Lueders, K. K. & Cullen, B. R. APOBEC3A and APOBEC3B are potent inhibitors of LTR-retrotransposon function in human cells. Nucleic Acids Res. (2006) doi:10.1093/nar/gkj416.

28. Jankowsky, J. L. et al. Mutant presenilins specifically elevate the levels of the 42 residue β-amyloid peptide in vivo: Evidence for augmentation of a 42-specific γ secretase. Human Molecular Genetics (2004) doi:10.1093/hmg/ddh019.

29. Ribet, D. et al. An infectious progenitor for the murine IAP retrotransposon: Emergence of an intracellular genetic parasite from an ancient retrovirus. Genome Res. (2008) doi:10.1101/gr.073486.107.

30. Welker, R., Janetzko, A. & Krausslich, H. G. Plasma membrane targeting of chimeric intracisternal A-type particle polyproteins leads to particle release and specific activation of the viral proteinase. J. Virol. (1997) doi:10.1128/jvi.71.7.5209-5217.1997.

31. Douville, R., Liu, J., Rothstein, J. & Nath, A. Identification of active loci of a human endogenous retrovirus in neurons of patients with amyotrophic lateral sclerosis. Ann. Neurol. 69, 141–151 (2011).

32. Tam, O. H. et al. Postmortem Cortex Samples Identify Distinct Molecular Subtypes of ALS: Retrotransposon Activation, Oxidative Stress, and Activated Glia. SSRN Electron. J. (2019) doi:10.2139/ssrn.3364349.

33. Schmitt, K. et al. Comprehensive Analysis of Human Endogenous Retrovirus Group HERV-W Locus Transcription in Multiple Sclerosis Brain Lesions by High-Throughput Amplicon Sequencing. J. Virol. (2013) doi:10.1128/jvi.02388-13.

34. Kassiotis, G. Endogenous Retroviruses and the Development of Cancer. J. Immunol. (2014) doi:10.4049/jimmunol.1302972.

35. Li, W. et al. Human endogenous retrovirus-K contributes to motor neuron disease. Sci. Transl. Med. 7, 307ra153 (2015).

36. De Cecco, M. et al. L1 drives IFN in senescent cells and promotes age-associated inflammation. Nature 566, 73–78 (2019).

37. Goodier, J. L. Restricting retrotransposons: A review. Mobile DNA (2016) doi:10.1186/s13100-016-0070-z.

38. Frost, B., Hemberg, M., Lewis, J. & Feany, M. B. Tau promotes neurodegeneration through global chromatin relaxation. Nat. Neurosci. 17, 357–366 (2014).

39. Mansuroglu, Z. et al. Loss of Tau protein affects the structure, transcription and repair of neuronal pericentromeric heterochromatin. Sci. Rep. (2016) doi:10.1038/srep33047.

40. Gold, J. et al. Safety and tolerability of Triumeq in amyotrophic lateral sclerosis: the Lighthouse trial. Amyotroph. Lateral Scler. Front. Degener. (2019) doi:10.1080/21678421.2019.1632899.

41. Bolger, A. M., Lohse, M. & Usadel, B. Trimmomatic: a flexible trimmer for Illumina sequence data. Bioinformatics 30, 2114–2120 (2014).

42. Andrews, S., Krueger, F., Seconds-Pichon, A., Biggins, F. & Wingett, S. FastQC. A quality control tool for high throughput sequence data. Babraham Bioinformatics. Babraham Institute (2015).

43. Dobin, A. et al. STAR: Ultrafast universal RNA-seq aligner. Bioinformatics (2013) doi:10.1093/bioinformatics/bts635.

44. Jin, Y., Tam, O. H., Paniagua, E. & Hammell, M. TEtranscripts: A package for including transposable elements in differential expression analysis of RNA-seq datasets. Bioinformatics (2015) doi:10.1093/bioinformatics/btv422.

45. Love, M. I., Huber, W. & Anders, S. Moderated estimation of fold change and dispersion for RNA-seq data with DESeq2. Genome Biol. 15, 550 (2014).

46. Ginestet, C. ggplot2: Elegant Graphics for Data Analysis. J. R. Stat. Soc. Ser. A (Statistics Soc. (2011) doi:10.1111/j.1467-985x.2010.00676_9.x.

47. Kolde, R. Package `pheatmap’. Bioconductor (2012).

48. DeVos, S. L. et al. Tau reduction in the presence of amyloid-β prevents tau pathology and neuronal death in vivo. Brain (2018) doi:10.1093/brain/awy117.

49. Altschul, S. F., Gish, W., Miller, W., Myers, E. W. & Lipman, D. J. Basic local alignment search tool. J. Mol. Biol. (1990) doi:10.1016/S0022-2836(05)80360-2.

50. Bao, W., Kojima, K. K. & Kohany, O. Repbase Update, a database of repetitive elements in eukaryotic genomes. Mob. DNA (2015) doi:10.1186/s13100-015-0041-9.

51. Benjamini, Y., Krieger, A. M. & Yekutieli, D. Adaptive linear step-up procedures that control the false discovery rate. Biometrika (2006) doi:10.1093/biomet/93.3.491.

52. Wickham, H. *ggplot2*. ggplot2 (2009). doi:10.1007/978-0-387-98141-3.

